# Modelling plastics exposure for the marine biota: Risk maps for fin whales in the Pelagos Sanctuary (North-Western Mediterranean)

**DOI:** 10.1101/538058

**Authors:** Federica Guerrini, Lorenzo Mari, Renato Casagrandi

## Abstract

The Mediterranean basin is among the most impacted marine ecoregions globally, being at the same time semi-enclosed by densely populated countries and crossed by trafficked maritime routes. Such anthropogenic pressure threatens both the quality of its waters and the high biodiversity living in them, making the role of Marine Protected Areas (MPAs) crucial for preserving species suitable habitats. Under the European Union Marine Strategy Framework Directive, marine litter has been recognized as one of the principal causes of marine pollution, and public awareness on its environmental and biological impacts is raising. Using a quantitative and data-driven modelling approach, here we assess the presence of plastic waste within the feeding grounds of the fin whale *Balaenoptera physalus*, an endangered cetacean for which there is increasing evidence of impacts due to microplastic ingestion. To this end, we analyze a decade (2000 - 2010) of advection patterns of marine plastic litter, modelled with a Lagrangian approach. Particles are released in the MPA Pelagos, the International Sanctuary for the Protection of Mediterranean Marine Mammals (North-Western Mediterranean, between France, Italy and Monaco), from different sources (i.e., untreated waste along coasts, plastic discharged from rivers and plastic pollution released along maritime shipping routes). Risk of exposure of fin whales to microplastic pollution is evaluated by interlacing plastic litter distribution maps obtained through modelling with maps of suitable habitats obtained from the elaboration of satellite chlorophyll-a data in species-specific visited areas. Our modelling results show that all the three main sources of plastic litter taken into account clearly contribute to impacting cetaceans in the Sanctuary, yet in a different manner. The procedure formalized here can be extended to assess the risk caused by plastic pollution in other MPAs as well as to evaluate possible impacts on other taxa, thus informing targeted actions to tackle the complex issue of marine litter.

## Introduction

Plastic materials have undisputably revolutionized our daily life, however their countless purposes are reflected by their ubiquitous presence as litter in the environment. The problem of plastic pollution and its impacts, in particular on marine ecosystems, have been known for decades (1,2), and interactions with marine biota have been observed since the 1950’s (3,4), less than ten years after the discovery of the most used polymers. In recent years, several accumulation and retention areas for plastic debris have been detected at the locations of the five main subtropical oceanic gyres at the global scale (5-8). These areas, frequently referred to as *garbage patches*, are showing increasing litter concentrations (9) due to the coupling between rising plastic production (10) and the input of plastic waste in the world’s oceans (11).

The European Marine Strategy Framework Directive (2008/56/EC, Descriptor 10), launched in 2008 with the main goal of achieving Good Environmental Status of European marine waters by 2020, recognized marine litter as one of the main causes of marine pollution. The Mediterranean Sea is no stranger to the issue of marine litter: some samplings and modelling experiments revealed concentrations similar to those found in the North Atlantic Gyre (7,12). The Mediter-ranean basin is, in fact, among the most impacted ecoregions globally due to increasing anthropogenic pressure (13), caused by rising coastal populations (14) and the intensification of maritime traffic (15), worsened and amplified by the introduction of alien, invasive species (16-18) and by the impacts of climate change (19). At the same time, the Mediterranean Sea hosts about 7% of the world’s marine biodiversity, with about 17,000 marine species (20) despite its relatively small size, as low as 0.32% of the whole ocean water content (21). At least 134 species living in the Mediterranean Sea have already been found to be affected by floating or seafloor litter, including some species of commercial value (22) and other endangered ones, like some cetaceans (23).

The effects and long-term consequences of plastic pollution on marine ecosystems, ranging from harm on wildlife due to entanglement or ingestion (22), biomagnification (24) and leaching of plastic additives (25), are causing concern at the global scale. Modelling experiments have been regarded as a useful tool to address the knowledge gaps about sources, pathways and accumulation areas of plastic, complementary to on-field sampling data (26). The use of models may be particularly effective in the Mediterranean Sea, where marine litter does not accumulate in garbage patches due to seasonal variations in surface ocean circulation patterns (27,28), thus possibly resulting in more complex spatio-temporal patterns. As pointed out by (12), litter dynamics in the Mediterranean basin resemble more *a plastic soup*. For this reason, sampling campaigns are key to provide a temporal snapshot of the concentrations of plastic items, but might not be representative of their long-term distribution. Ocean circulation models and oceanographic reanalyses are commonly applied when studying marine litter distribution both for the global oceans (6) and for the Mediterranean Sea (28-30). They allow us to account not only for the transport of particles within ecologically relevant seasons, but also for its inter-annual variability. Moreover, oceanographic modelling can provide some insight into the exposure of marine biota to plastic litter, by coupling simulation outputs with ecological information (such as maps of species distribution) which can be prodromal to more in-depth ecotoxicological studies investigating species-specific dose-response impacts of plastic pollution (31). At present there are only few examples along this line of research. (32) evaluated the risk of entanglement in derelict fishing gear for sea turtles in the nearshore waters of Northern Australia. To do so, they elaborated the spatial distribution of drifting nets, obtained from both Lagrangian numerical models and beach cleanup data on the oceanographic side, and turtle bycatch records on the ecological side, as gathered from fishing activities in the area. A similar approach was followed to assess the risk at global scale of plastic ingestion by seabirds (33) and sea turtles (34). In these two later studies, the risk was defined as the probability of plastic ingestion, predicted by applying a logistic regression model on some biologically relevant characteristics of stranded individuals (such as life-history traits and body size). Some recent items of research targeted specifically the Mediterranean Sea. (35) evaluated the exposure of sea turtles to marine litter with aerial surveys covering the French Mediterranean and Atlantic waters, identifying copresence, encounter probability and density of debris surrounding individuals. (36) were first in proposing a qualitative comparison method for visually identifying regions endowed with high risk of exposure to plastics. They contrasted 15 days of litter distribution from oceanographic modelling and areas with high potential habitat value for a target species, the feeding grounds of the fin whale *Balaenoptera physalus* in Pelagos Sanctuary.

In this work, we combined a decadal Lagrangian particle tracking with species-specific habitat suitability maps to obtain a quantitative, georeferenced risk indicator for the marine ecosystem of the Pelagos Sanctuary, the largest Marine Protected Area of the Mediterranean Sea located in its North-Western side. To this aim, we exploited a simplified version of the habitat suitability model calibrated by (37) targeted on the fin whale *B. physalus*, an endangered cetacean species which aggregates in Pelagos during the summer season for feeding (38,39) and for which there is increasing evidence of microplastic ingestion in its foraging area (40). Improving over (36), we formalize a procedure to obtain multi-annual risk maps for fin whales in the Pelagos Sanctuary, an approach that can potentially be extended to other marine areas and/or biota.

## Methods

As typically done in risk assessment studies (41), we obtained the *risk* associated with plastic pollution for the marine biota as the product of a *hazard* factor and an *exposure* factor. This definition required us to follow three sequential steps during this work, one for the evaluation of each factor: 1)the hazard, determined by plastic densities as simulated using oceanographic modelling over the Pelagos Sanctuary domain; 2) the exposure, obtained by mapping habitat suitability for the fin whale; and 3) the risk, computed by multiplying cell by cell the outcomes of the hazard and exposure steps.

### A. The hazard — maps of plastic density

Plastic density has been modelled by simulating the advection of Lagrangian particles using oceanographic reanalyses produced by the Copernicus Marine Environment Monitoring Service (42). We used surface currents data (zonal and meridional components), while we neglected vertical velocities (Figure 1A). The spatial extent of our simulation domain (3.5 °E to 12.5°E, 38.5°N to 4°N on a 1/16°x 1/16°oceanographic grid) is wider than the Pelagos Marine Protected Area for a more realistic simulation of particle transport on a larger scale. In fact, litter may enter the focal area from sources that are located outside the Sanctuary, or temporarily leave the Sanctuary waters before eventually re-entering. We located several release points along the coastlines, at the mouths of the major rivers and along the main maritime routes to mimic real-life sources of plastic litter, as it will be detailed below. Simulations have been run, with a daily step, in order to cover the summer seasons of the years 2000-2010. This choice is based on the ecology of the fin whale, as mentioned above, and it is coherent with the sightings dataset used by (37) to calibrate their suitability model. To allow model warm-up before collecting simulation outputs in the summer seasons, we begun each year of simulation in March and terminated it in September. Removal of plastic fragments from the surface layer has been included in the model by characterizing Lagrangian particles with a certain source-independent permanence period in the surface layer, corresponding in fact to the duration of their transport. Assuming that particle removal is a Poissonian process, the permanence period of each particle can be extracted from an exponential distribution with an average decay rate of 50 d^*−*1^(see 30). The distribution of the permanence times was then approximately discretized into three classes with relative frequency of 1/3 each. We thus divided the particles released every day from each site into three groups, with short (10 days), medium (36 days) and long (105 days) duration of transport before removal from the simulation. The calculated advection times are consistent with the existing literature (43-45). However, since the model is effectively memoryless after 105 days, the effects of those (rare) particles with longer residence times in the surface layer might be underestimated.

**Fig. 1.**
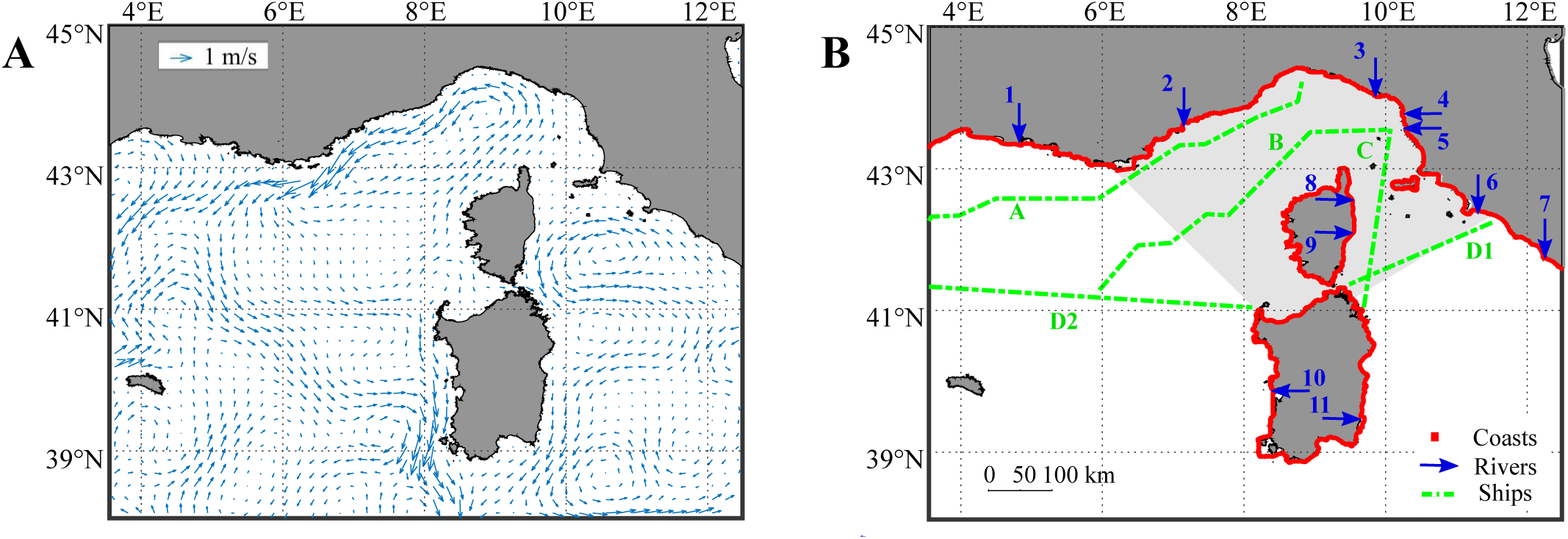
Input data used to assess the hazard component of the risk maps via numerical simulations. **(A)** Surface circulation fields in the simulated summer seasons (averaged over the time horizon 2000-2010), as obtained from reanalysis data. **(B)** Geographic displacement of particle sources used in the simulations. The Pelagos Sanctuary is shaded in gray. Coastline release sites are colored in red. Blue arrows indicate the mouths of the major rivers flowing directly into the simulation domain and have been numbered from West to East: 1. Rhone, 2. Var, 3. Magra, 4. Serchio, 5. Arno, 6. Ombrone, 7. Tevere, 8. Tavignano, 9. Golo, 10. Tirso, 11. Flumendosa. Green dash-dotted lines indicate the main shipping lanes: a. Genova-Barcelona, b. Livorno-Barcelona, c. Livorno-Olbia, Civitavecchia-Barcelona eastern (d1.) and western (d2.) sections.

Particle release locations within the focal study area are shown in Figure 1B. Plastic release from coasts was uniformely distributed along 2,500 km of coastlines, with a spacing of approximately 650 m between each source point, finally resulting in 3,843 release locations. Coastal areas acting as particle sources encompassed the Italian regions of Liguria, Tuscany (including the major island of its Archipelago, Elba island), Sardinia and part of Lazio, and the French Cote d’Azur, part of the Languedoc-Roussillon coast and Corsica. Plastic discharged from rivers was assumed to come from the outflows in the relevant area of the eleven major rivers, i.e. those characterized by the highest yearly average discharges. The rationale behind our discharge-driven choice criterion lies in the positive correlation between riverine discharge and the amount of plastic released by a river, as pointed out by several studies (see e.g. 46, 47). Plastic litter produced by maritime activities was accounted for by modelling plastic sources along the main naval routes (commercial and transport/touristic) with a 2 km spacing between source points. We extrapolated the most trafficked routes from the SafeMED GIS (48) and (15).

From a technical perspective, there is a trade-off between the number of particles per source point to be used for simulation (number that should be maximized so as to have a homogeneous coverage of the domain while computing the hazard) and the computational load. After a testing phase, we chose to differentiate the number of released particles for each source type. A good compromise was found by releasing every day 300 particles from each coastal and maritime source and 30,000 from each river. The initial position of each particle was randomly extracted within a 100 m radius from its source point, to increase spatial variability and somehow mimic occasional release. Moreover, the vertical position of the particles was randomly assigned within the surface layer of the oceanographic circulation reanalyses (0 - 1.47 m) and kept fixed thereafter. Overall, considering all source types, a total of 1,739,100 particles was released every day, amounting to about 3 billion particles tracked over an 11-year-long time window.

The contribution to accumulation of plastics from each of the three source types has been simulated considering them independently. Then, to obtain an aggregated picture of plastic within the study area, daily outputs of the single-source simulations have been normalized by dividing the number of particles in each cell of the oceanographic grid by the maximum number of particles registered in all cells. To account for the different magnitude of the plastic input from coastlines, rivers and shipping routes, the aggregated mapping was obtained by attributing different weights to the particles according to their source, with the coast-to-rivers-to-shipping lanes ratio of 50%:30%:20% proposed by (30).

In terms of the temporal dimension of particles concentration, at first the simulation outputs from the three source types were averaged over the eleven summer seasons simulated. Then, inter-annual variability was estimated for each source type by considering, summer by summer, the fraction of particles that entered the Pelagos Sanctuary on the total amount released in a year of single-source simulation. Annual fluctuations of plastic distribution within the Sanctuary was quantified through the coefficient of variation (CV), computed on the aggregated particle density maps as a cell-by-cell ratio between standard deviation and mean.

### B. The exposure – maps of habitat suitability

To specifically target the exposure of the fin whale in our analyses, we determined habitat suitability for the species by applying a simplified version of the model proposed by (37). In their niche model, Druon and collaborators used a multi-criteria evaluation to define the main features of the whales’ habitat, providing a specific chlorophyll-a (chl-a) range, a minimum threshold for the norm of the horizontal gradient of sea surface temperature and chl-a, and a minimum water depth. Among these predictors, we selected the range of chl-a concentration (0.11 to 0.39 mg/m^3^) and water depth (from 200 to 2800 m) to identify potential habitats. Our simplification is supported by two considerations: in the mentioned study, chl-a fronts were found to be the main predictor of whale presence, and most (82%) of the recurrent potential habitat identified by (37) has been identified in correspondence of those water depths. To determine sea surface chl-a concentration, we used monthly satellite data from MODIS-Aqua, the Moderate Resolution Imaging Spectroradiometer aboard the NASA-Aqua satellite, obtaining georeferenced data such as those shown in Figure 2A. Data were available from 2002 to 2010, since chl-a measurements started in late June 2002. Water depth was taken into account by using the General Bathymetric Chart of the Oceans (49; Figure 2B). We thus obtained the fin whale suitable habitat by assigning value 1 to the cells of the oceanographic grid which matched our criteria and 0 to the others, thus obtaining the exposure factor as Boolean masks for the summer months of the years 2002-2010.

**Fig. 2.**
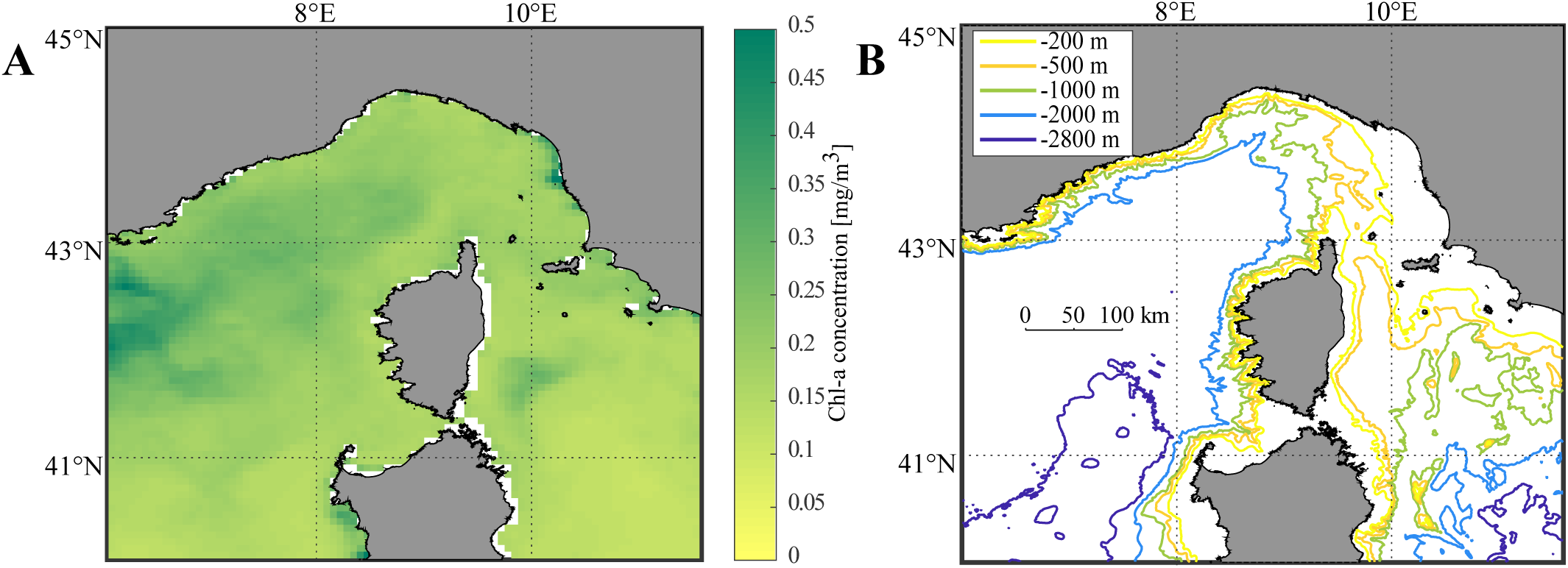
Input data used to assess the exposure component of the risk maps using habitat suitability modelling. **(A)** An exemplification of a temporal snapshot (June 2006) of surface chlorophyll-a (chl-a) concentration satellite data, see text for data sources. **(B)** Bathymetry of the Pelagos Sanctuary area as provided by (49).

Besides fin whales, other species can be studied by selecting a relevant indicator of their habitat. As a notable example, the presence of higher trophic marine biota can be predicted using the Net Primary Production (NPP). Typically, areas with high NPP are in fact positively associated with abundance of phytoplancton, which is at the basis of the food chain, making reasonable to assume NPP as a proxy of ecosystem size (50).

### C. The risk – interlacing hazard and exposure

The risk associated with exposure to plastic for the marine taxa under study was finally calculated at a monthly temporal step, as the product between the two gridded fields described above: the monthly-averaged plastic density data (hazard factor, as resulting from our Lagrangian simulations) and the monthly masks of fin whale potential suitable habitat (exposure factor), that is:

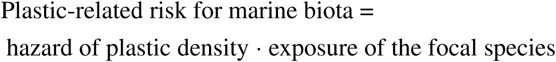

As a consequence of MODIS-Aqua data availability as mentioned above, we could calculate risk maps for the fin whale from July 2002 and not before.

## Results

Figure 3 summarizes the contribution to plastic density in the area of the Pelagos Marine Protected Area (MPA) from each of the three sources – coastlines (panel A), main shipping routes (panel B) and major rivers (panel C). At a glance, release of particles from coastlines and maritime traffic (Figures 3A and 3B) present comparable particle distributions, in terms of both the coverage of the simulation domain and of geographical patterns that can be identified. For example, the high-density areas located in the Central Ligurian Sea, in the Gulf of Lion and a lesser one in the Tyrrhenian Sea emerge from both single-source simulation settings. Particle dispersion from these two linear sources appears to be more homogeneous than the one generated by rivers (Figure 3C). At the geographical scale of interest, rivers behave in fact like punctual sources, releasing at their mouth a plume of particles whose dispersion depends on the dynamicity of the waters they flow into. In the riverine setting, the Northern Tyrrhenian Sea presents higher particle concentration than the Ligurian Sea.

**Fig. 3.**
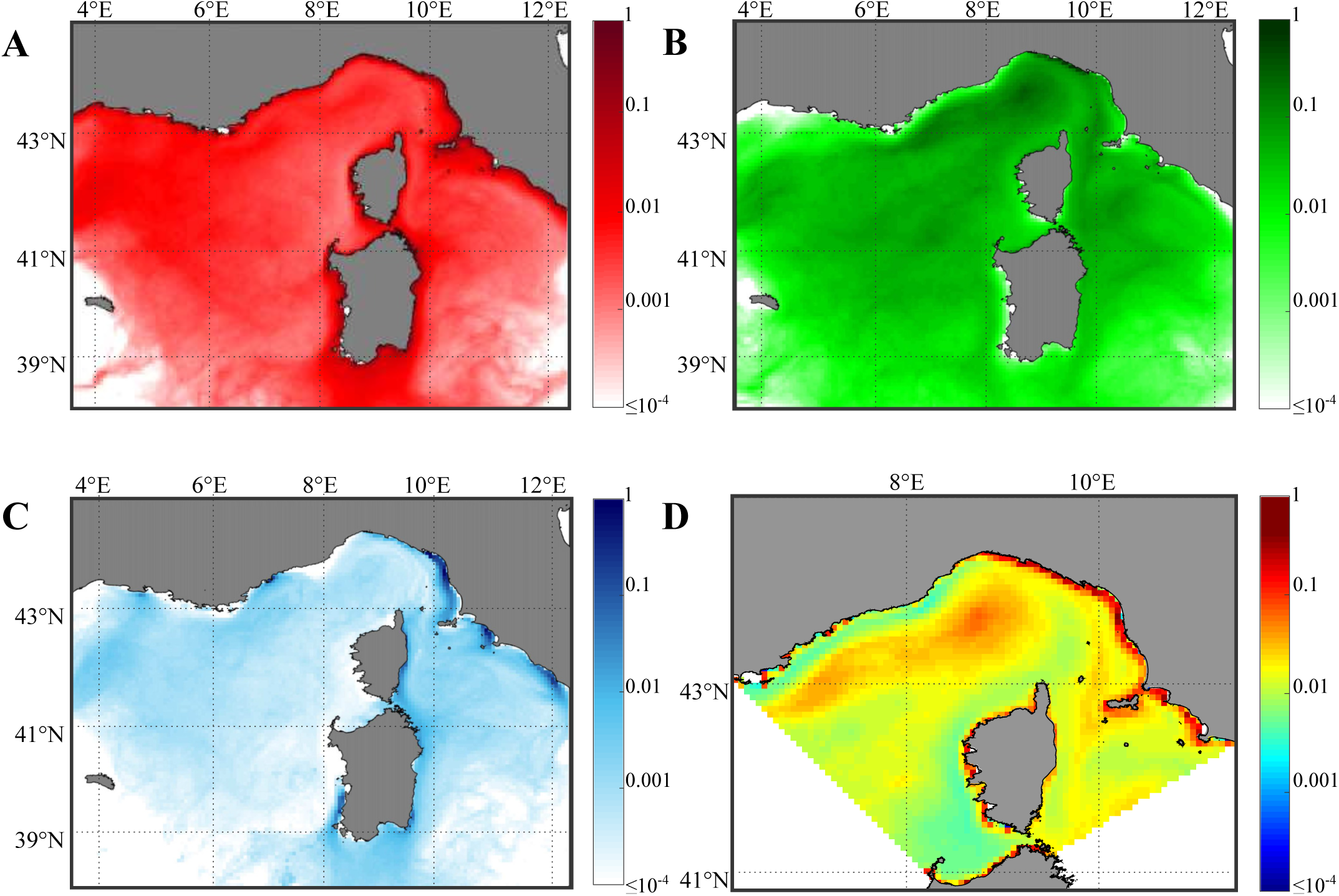
Maps of summer plastic density (hazard component) averaged over the period 2000-2010 from each source of release: **(A)** coasts, **(B)** ships and **(C)** rivers. **(D)** Overall decennial hazard map in the Pelagos Sanctuary, obtained with a weighted aggregation.

Particle concentration hotspots emerge more clearly when considering how the three pollution sources act all together (Figure 3D). Again, particle density is high in the Central Ligurian Sea, forming a plume-like structure that extends westwards, starting from offshore the Ligurian coasts and reaching the French part of the domain. Interestingly, other hotspots that did not emerge in the single-source panels can now be clearly identified thanks to the weighting and aggregation procedure. One of those hotspots is located along the coastal marine areas of Tuscany and Lazio, on the eastern part of the domain, while another is a small offshore region between Tuscany and Corsica, on the western side of Elba island. It is worth noting that this last hotspot, despite being less pronounced than the other two, is already known to be a potential accumulation area for marine litter (12,36), possibly because of its proximity to a seasonal anticyclonic eddy structure named *Capraia gyre* (51) after the Capraia island, located north of Elba in the Tuscan Archipelago.

Mapping the average distribution of plastic provides information about the location and the extension of high concentration areas, however possible yearly fluctuations might not be visible. For this reason, in Figure 4 we show our two indicators of inter-annual variability. In Figure 4A, fluctuations in the amount of particles entering Pelagos are sharper for the release from rivers than from coastlines and shipping routes. This is probably due to the number and the geographic location of release sites included in each source type. In fact, coastlines and shipping routes are modelled as hundreds of evenly spaced point sources spread on a wide area, thus they are less affected by the variability of surface circulation in delivering particles to Pelagos. Conversely, dispersion of particles from rivers, which are fewer in number and described as point sources, can be more easily influenced by the actual circulation regime. In some years, surface currents appear to be particularly favourable in reducing the accumulation of riverine particles in Pelagos (e.g. 2002 and 2007, see again Figure 4A). Moreover, the two minima and the overall shape of the riverine time series in Figure 4A suggest a non-erratic, multiannual pattern. It can also be observed that the time series of plastic contributions to Pelagos from rivers and coastlines appear to be in phase, while the one associated with shipping routes is out of sync, even in counterphase in some years. The CV map shown in Figure 4B suggests that particle density is subject to sharper inter-annual variability in open seas rather than in coastal waters, as somehow expected since the latter are less exposed to strong currents. Furthermore, the hotspot linked to the Capraia gyre (described above) presents rather low variability, perhaps strengthening the evidence that this high-concentration area tends to build up frequently.

**Fig. 4.**
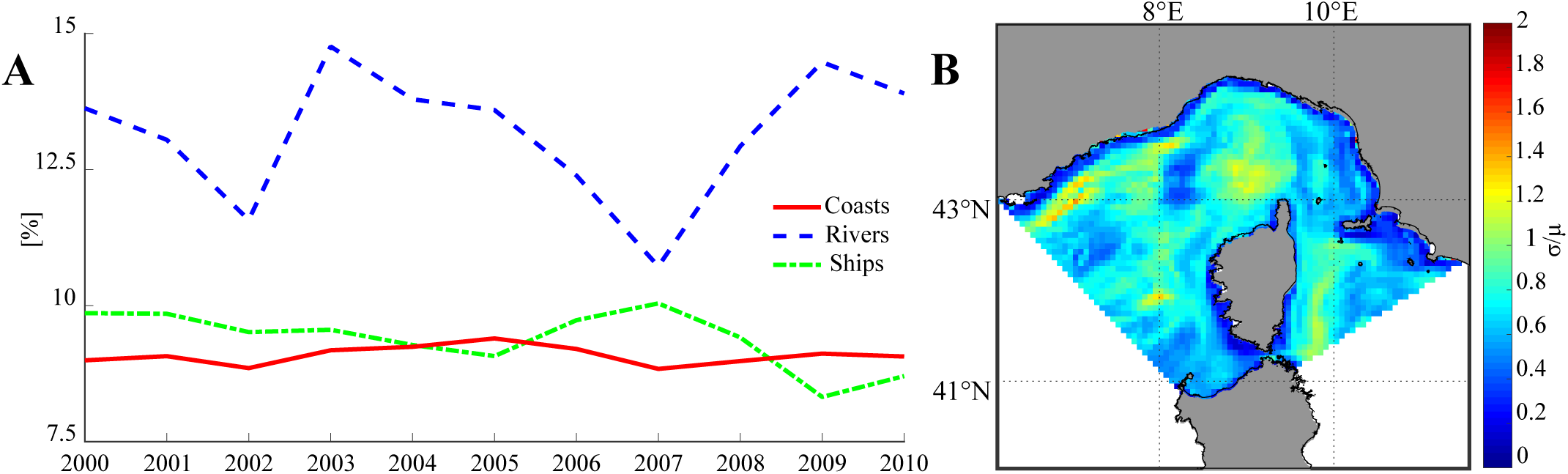
Indicators of inter-annual variability in plastic distribution (hazard component). **(A)** Partitioning by source type of particles retained in the Pelagos Sanctuary each year during the simulated period. Percentages refer to the total number of particles released in a year from the relevant type of source. **(B)** Inter-annual Coefficient of variation of plastic density over the Sanctuary area in 2000-2010.

The time-averaged exposure factor of fin whales in the Pelagos MPA (2002-2010) is shown in Figure 5A. Overall, the potential suitable habitat appears to cover the whole region, excluding the areas whose water depth did not match our suitability criteria. This result is not surprising at all, since Pelagos is well known for its highly productive waters and frequent whale sightings, being in fact a Sanctuary for cetaceans. Figure 5B finally integrates the hazard and the exposure monthly maps, and show the average risk for the period 2002-2010. On average, the risk for the target species appears to be maximum along the Ligurian and Western Corsica coasts. The widest hotspots match, comprehensibly, the high particle concentration areas previously identified. Again, the Ligurian sea presents an extended area with high risk, whose severity decreases towards the French coasts. In the eastern-most part of the domain, the risk appears to be lower than in the Ligurian hotspot. However, noticeable values can still be observed in correspondence of the Northern Tyrrhenian area and close to the Capraia gyre, the latter being less defined.

**Fig. 5.**
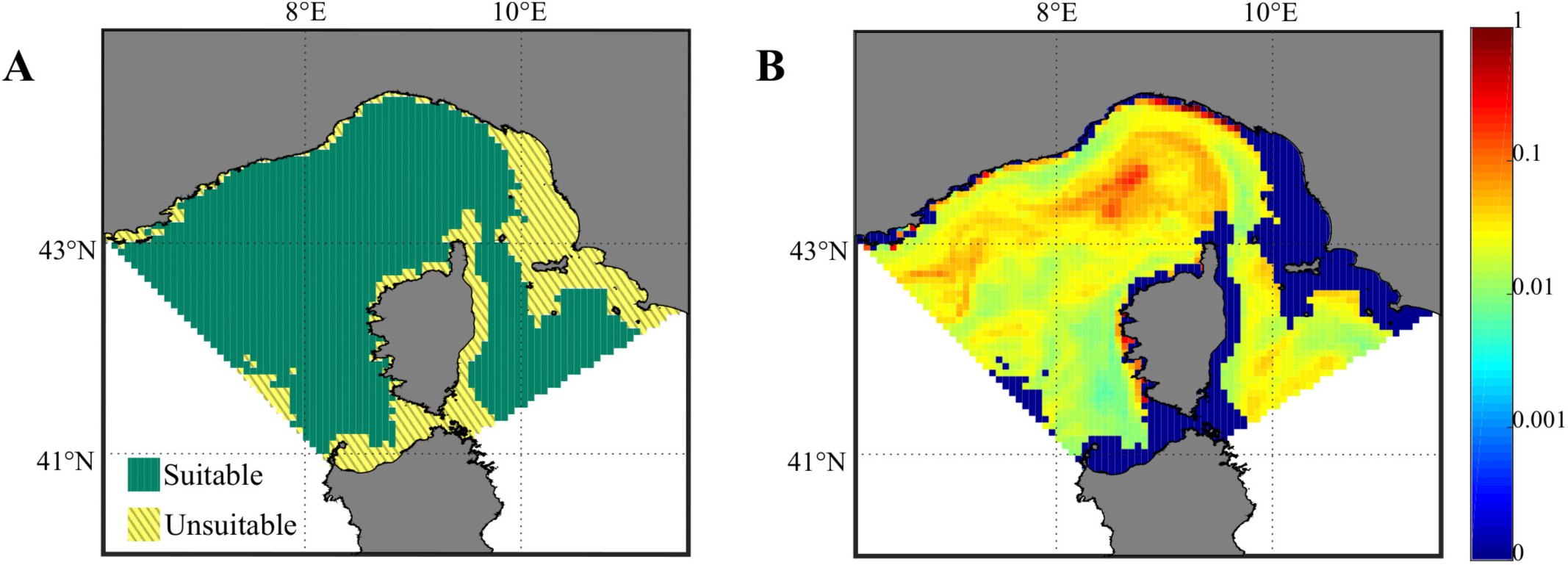
Maps of exposure to plastic and of the related risk for the fin whale. **(A)** Boolean mask representing the average suitable habitat in 2002-2010, calibrated using a simplification of the model by (37). **(B)** Average risk for *B. physalus* in 2002-2010, as resulting from multiplying the hazard and the exposure monthly maps.

## Discussion

In the present work, extensive Lagrangian simulations were run to model the surface advection patterns of plastic litter on a wide geographical domain embracing the Pelagos International Sanctuary for the Protection of Mediterranean Marine Mammals. Aim of our modelling study was to assess the potential distribution of plastic waste within the feeding grounds of the fin whale, an endangered cetacean inhabiting the Mediterranean Sea. Nearly three billion particles were released from coastlines, major rivers and most congested shipping lanes during the study period 2000-2010, and followed daily during their transport driven by surface ocean currents from the most up-to-date oceanographic reanalyses. In accordance with existing literature, we assumed that particle transport duration could be taken from an exponential distribution with average decay rate of 50 days. We modelled the fin whale suitable habitat on satellite-derived and bathymetry data, using a selected subset of the criteria identified by (37) based on whale sightings occurred during the decade 2000-2010. The suitable areas detected as exposure component were used to finally compute the risk related to plastic pollution for the fin whale, that is proportional to the hazard of finding particles in the region.

Over the ecologically relevant summer months of the decade 2000-2010, our simulations show that the highest average particle densities were localized in the Ligurian Sea, in the Northern Tyrrhenian Sea and between the Tuscan Archipelago and Corsica, as well as along the eastern coastlines of the simulation domain. What stands out in every examined plot is that no area within the Pelagos Sanctuary appears to be unaffected by the potential pollution sources that have been selected in this work. Our modelling results are in a rather good agreement with the observational data of plastic pollution based on sampling procedures by (12), and (52), except for a high-density area that the latter found in the Bonifacio strait, not reproduced by the present elaborations. We note that this region is relatively poorly resolved in the oceanographic circulation reanalyses we exploited. Our results are also comparable with the simulation results by (7), (36) and (30), while there are some dissimilarities with the work by (28), probably because of the use of different oceanographic grids. In addition, our analyses confirm a known feature of the summer oceanography of the Pelagos area: the Capraia gyre. Simulations show that this region is also endowed with a low coefficient of variation, underlying its recurrent and persistent appearance in the summer seasons of the years considered.

The potential whale habitat we modelled here, visible in Figure 5A, is similar to both the one obtained by (37) and the fin whale presence probability observed by (53). Risk from floating microplastic debris to the endangered *B. physalus* was obtained by properly interesecting the elaboration of the particle density patterns (the hazard) and indicators of potential habitat suitability for this species (the exposure). Our method highlighted a potential risk hotspot for the fin whale in the Liguro-Provençal basin, which is coherent with the overlay of plastic distribution and potential habitat qualitatively identified and discussed by (36). This area notoriously plays a crucial role for the species, especially during its summer feeding season (37,39), thus the high risk of plastic pollution we found there may be of great concern.

Our study was focused on a particular species on a key MPA, the largest of the Mediterranean Sea. However, the proposed approach can be potentially extended to other marine species to assess whether they are threatened by plastic pollution and to which extent, as well as to other areas of interest. A practical example of such extension of our approach is shown in Figure 6, where we applied the procedure hereby explained by using as exposure factor the monthly NPP maps for the Pelagos Sanctuary area (from the Mediterranean Sea Biogeochemistry Reanalyses available from the Copernicus Marine Environment Monitoring Service; 54). Although the effects of microplastics on pelagic productivity are still uncertain at current environmental concentration levels (55), NPP can be used to assess the exposure of the marine ecosystem (50), as it is linked to phytoplancton growth, a main source of nutrition for higher trophic levels.

**Fig. 6.**
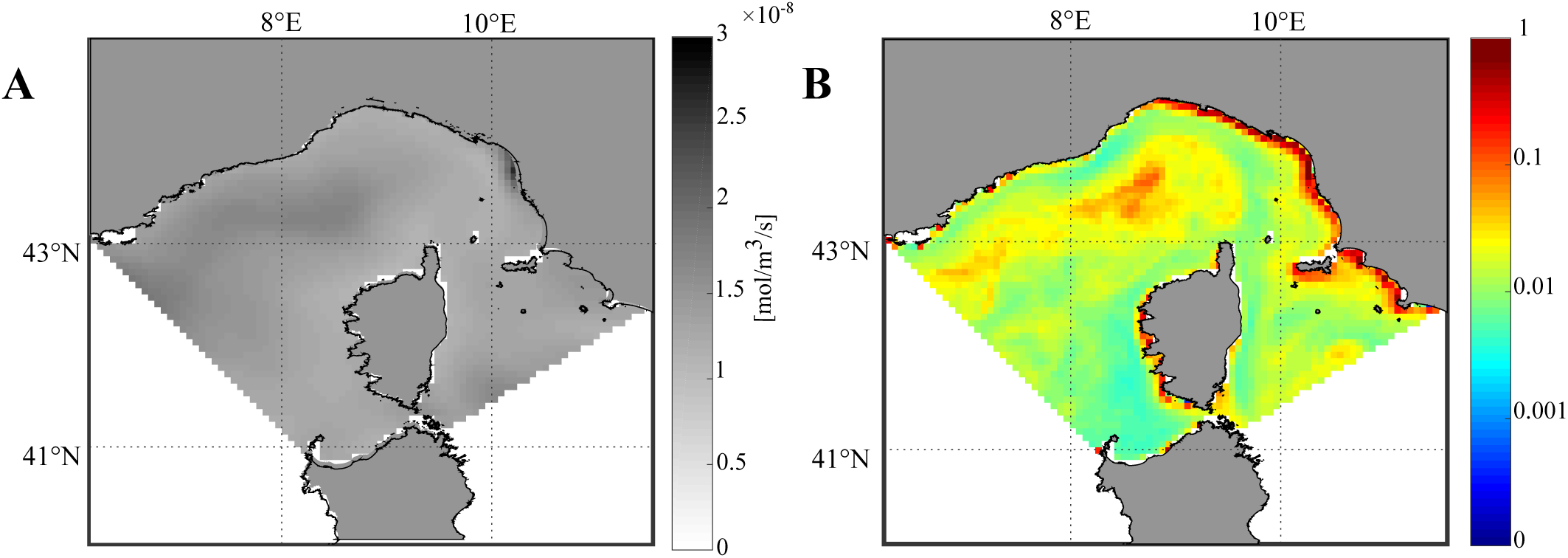
An application of our methodology to a different ecological target: maps of exposure to plastic pollution and of the related risk for the whole marine ecosystem of the Pelagos Sanctuary during summer months. **(A)** Average NPP in 2000-2010, as derived from (54), used to assess the exposure component for the marine ecosystem. **(B)** Average risk caused by plastic pollution.

To conclude, we performed a first decadal assessment of plastic-related risk for the fin whale in the Pelagos MPA supported by both numerical simulations and ecological modelling, providing a simple methodology to obtain a quantitative, species-specific risk indicator. Improving over other existing risk assessments along the same line of research, modelled particles were released from the major sources, aiming to simulate real-life plastic input. Although being promising, the method we propose here would significantly benefit from a more precise characterization of plastic inputs from the relevant sources, both in terms of masses and sizes of the items. Further research needs to be carried out in this direction, also to better define seasonal variability of the sources, e.g. due to tourism or river runoff. Additionally, little is known yet about the residence time of floating plastics on the sea surface. A better knowledge of the behaviour of plastic waste in the marine environment would improve not only the modelling of its dispersion patterns but also the quantification of the standing stocks in the different compartments (ocean surface, seafloor and beaches; 56). Combining these sources of information can be helpful to address both effective waste management policies and volunteering actions (like clean-ups), aimed to reduce inputs. Tackling plastic pollution is a worldwide challenge with important socio-economic and environmental implications at all scales (from local to national and beyond), involving a variety of stakeholders. As the primary solution to the problem appears to be preventing contamination in the first place (57), scientifically-informed, targeted actions are nowadays needed more than ever.

## ACKNOWLEDGEMENTS

We acknowledge funding from the H2020 project “ECOPOTENTIAL: Improving future ecosystem benefts through Earth observations” (grant agreement No. 641762). We are also thankful to Arianna Azzellino for feedbacks on our study and to Gaetano Leoni for stimulating discussions.

